# Amyloid conformers of the FXR1 protein prevent mRNA degradation in cortical neurons

**DOI:** 10.1101/544650

**Authors:** J.V. Sopova, E. I Koshel, T.A. Belashova, S.P. Zadorsky, A.V. Sergeeva, V.A. Siniukova, A.A. Shenfeld, M.E. Velizhanina, K.V. Volkov, A.A. Nizhnikov, E.R. Gaginskaya, A.P. Galkin

**Author notes:** Current address: Department Chemistry and Molecular Biology, ITMO University, 191002 St. Petersburg, Russian Federation. Current address All-Russian Research Institute for Agricultural Microbiology, Pushkin, St. Petersburg 196608, Russian Federation. **Corresponding Author**: A.P. Galkin, Vavilov Institute of General Genetics, St. Petersburg Branch, Russian Academy of Sciences, 199034 St. Petersburg, Russian Federation.

## Abstract

Functional amyloids regulate vital processes in a variety of organisms from bacteria to higher eukaryotes. The development of methods enabling large-scale screening for amyloids opens up opportunity for systemic analysis of the prevalence of amyloids in nature. Using an original proteomic approach, we identified several proteins forming amyloid-like detergent-resistant aggregates in the rat brain. One of them is the FXR1 protein, which is known to regulate memory and emotions (1, 2). We demonstrated that in brain FXR1 forms amyloid oligomers and insoluble detergent-resistant aggregates that strongly colocalize with amyloid-specific dye Thioflavin S and bind mRNA molecules. Moreover, we demonstrated that mRNAs colocalized with FXR1 amyloid particles are completely resistant to treatment with RNAse A. Taking into consideration that the members of ribonuclease A superfamily function in neurons (3) we can conclude that amyloid conformers of FXR1 control RNA stability in brain. Thus, in contrast to pathological amyloids that cause neurodegeneration, FXR1 is the functional amyloid in forebrain. We showed that amyloid properties of FXR1 depend on its N-terminal part from 1 to 379 amino acids. This fragment forms amyloid fibrils *in vitro* that bind Congo red and manifest apple-green birefringence when assayed by polarization microscopy. The amyloid-forming region of FXR1 is highly conserved in mammals. These data suggest that the ability of amyloid conformers of FXR1 to protect mRNAs is characteristic of different mammalian species, including humans.

**Significance Statement:** Amyloids are highly ordered cross-β sheet protein fibrils associated with many neurodegenerative diseases including Alzheimer’s disease. However, some amyloid proteins regulate vital processes. We identified a set of proteins that form amyloid-like aggregates in the brain of healthy rats. One of them - the FXR1 protein is known to regulate memory and emotions. FXR1 forms amyloid fibrils that bind RNA molecules and prevent their degradation in brain cortex neurons. Amyloid-forming sequence of FXR1 is highly conserved across mammals including human. Discovery of functional amyloids in mammalian brain shows that strategy aimed at the development of universal anti-amyloid drugs is unpromising. Such potential drugs should prevent or suppress formation of pathological aggregates of a certain protein, but not affect functional amyloids.

## Introduction

Amyloids are non-branching fibrils that are composed of stacked monomers stabilized by intermolecular β-sheets arranged perpendicular to fibril axis. Such extra- and intracellular fibrils formed by various proteins were discovered in bacteria and eukaryotes (4, 5). Some proteins form pathological amyloid fibrils that cause neurodegenerative and other incurable human diseases, called amyloidoses, that affect millions of people worldwide (5). Other amyloids regulate vital processes. Functional amyloids in bacteria contribute to biofilm development and can generate toxic oligomers that cause damage to lipid membranes (6). A repeat domain of human Pmel17 protein forms fibrils essential for melanin deposition and synthesis in human pigment-specific cells (7). It was shown that mammalian neuropeptides and protein hormones are stored in an amyloid-like cross β-sheet-rich conformation in the pituitary secretory granules (8). There is reason to assume that functional amyloids can play a role in the regulation of memory. For example, proteins of the CPEB family form oligomers critical for the persistence of long-term synaptic facilitation and long-term memory in the brain of *Aplysia californica, Drosophila melanogaster and Mus musculus*. These oligomers, like classical amyloids, are resistant to treatment with sodium dodecyl sulfate (SDS) (9–11). However, SDS-resistance is a typical feature not only of amyloids, but also of some other filaments and protein complexes of a non-amyloid nature (12, 13). Besides, oligomers of CPEB bind amyloid-specific dyes only *in vitro* or when overexpressed *in vivo* but not under native conditions (9). Thus, the amyloid nature of the CPEB proteins under native conditions remains controversial.

Discovery of each new functional amyloid is a notable scientific event because until recently there were no methods for large-scale screening for amyloids. Recent advances in the development of a methodology of proteomic screening for amyloids allow to move from identifying individual amyloid proteins to systemic analysis of the prevalence and significance of amyloids in different species (12–15). These methods are based on the resistance of amyloid aggregates to treatment with SDS that makes it possible to separate them from most other non-amyloid protein complexes (16). The amyloid properties of the proteins identified in such screenings should be confirmed by further individual analysis. Here, we applied our original proteomic approach in order to search for functional amyloid-forming proteins in the rat brain. We identified several proteins that formed amyloid-like aggregates in brain and performed in-depth analysis of the amyloid properties of the FXR1 protein, which is involved in the regulation of memory and emotions (1, 2). We demonstrated that FXR1 forms amyloid conformers that bind RNA and prevent its degradation in brain cortex neurons. Comparative analysis of the amino acid sequences of FXR1 ortologs suggests that its amyloid properties are conserved at least in vertebrates.

## Results

### Proteomic screening identifies proteins forming detergent-resistant amyloid-like aggregates in the brain of *Rattus norvegicus*

To identify the proteins forming detergent-resistant amyloid-like aggregates in the brains of six-month-old males of *Rattus norvegicus*, protein lysate was obtained from the total brain samples. The fraction of proteins forming SDS-resistant aggregates was isolated and purified using the PSIA-LC-MALDI proteomic approach (12) optimized for brain tissues (see Materials and methods). The proteins present in this fraction were solubilized, treated with trypsin, separated by HPLC and identified by mass spectrometry. The experiment was repeated independently with brain samples obtained from three animals. The proteins NSF, MBP, RIMS1, STXB1 and FXR1 were identified by mass-spectrometry in all three analyzed samples (Table S1; Figures S1-S5). All of these proteins, except MBP, contain potentially amyloidogenic regions predicted by the ArchCandy algorithm (17) (Table S1). Notably, there are some data suggesting that the myelin basic protein (MBP), that is a component of specialized membrane covering axons, may form amyloid structures in the brain (18).

In this paper we focused on the analysis of the amyloid properties of the FXR1 protein that belongs to the Fragile X-Related (FXR) family of RNA-binding proteins, which also includes Fragile X Mental Retardation (FMRP) and Fragile X-Related 2 (FXR2) proteins (19). The FXR2 and FMRP proteins, in contrast to FXR1, were not identified in SDS-resistant fractions. The FXR1 protein identified in our screen binds specific RNAs and is involved in regulation of long-term memory, and emotion processing (1, 2). Taking into consideration these neurospecific functions of FXR1, we decided to comprehensively analyze its amyloid properties *in vivo* and *in vitro*.

### FXR1 forms functional amyloid conformers in brain that bind mRNA and prevent its degradation

To check whether FXR1 aggregates *in vivo*, total protein lysate from the rat brain was separated into three fractions: 1) proteins less than 100 kDa; 2) monomers and soluble oligomers larger than 100 kDa; 3) insoluble aggregates. Using an FXR1-specific antibody we demonstrated that this protein is predominantly present in fractions containing oligomers and insoluble aggregates (Figure 1A and 1B). Monomers of FXR1 were almost undetectable in the brain lysate (Figure 1A). Two bands are detected in the oligomeric fraction and four bands are found in the fraction containing insoluble aggregates (Figure 1A). These results are in complete agreement with previously published data, according to which several isoforms of FXR1 protein exist in the mouse and rat brain due to alternative splicing of 3’ part of FXR1 mRNA (20–22).

**Fig. 1.**
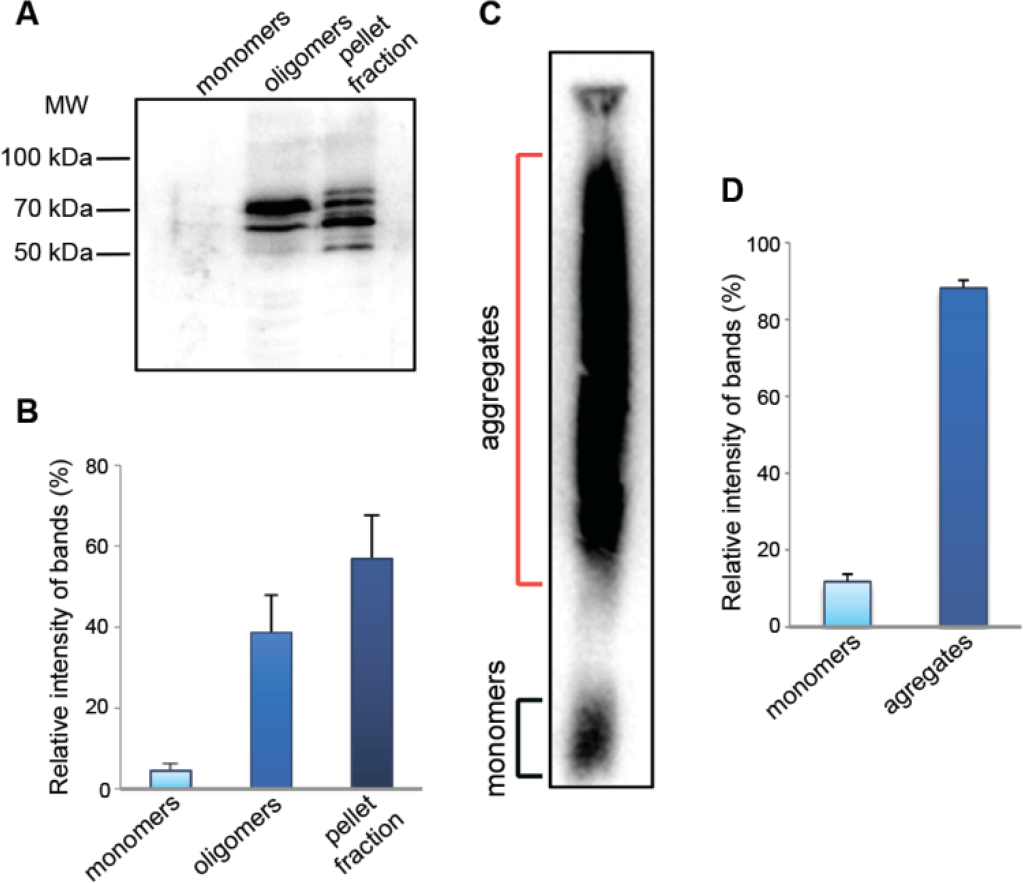
FXR1 forms SDS-resistant aggregates in rat brain. (A) FXR1 presents in brain in fraction of oligomers and insoluble aggregates. (B) Relative intensity of bands corresponding to FXR1 monomers, oligomers and insoluble aggregates is represented as mean ± SEM for three independent brain samples. (C) A large portion of FXR1 in rat brain forms SDS-resistant aggregates. (D) Relative intensity of bands corresponding to FXR1 monomers and SDS-resistant conformers is represented as mean ± SEM for three independent brain samples.

Known amyloids form SDS-resistant aggregates *in vivo* that may be detected by semi-denaturing detergent agarose gel electrophoresis (SDD-AGE) (23, 24). The total brain lysate was treated with 1% SDS and separated by agarose gel electrophoresis. A large portion of FXR1 formed detergent-insoluble aggregates (Figures 1C and 1D). This result resembles the data of proteomic screening for amyloid-forming proteins according to that FXR1 forms SDS-resistant aggregates *in vivo* in all rat brain samples analyzed (Figure S5).

To verify our data suggesting that FXR1 is present in amyloid form in brain, we compared the localization of FXR1 and amyloid-specific dye Thioflavin S on cryosections of the brain cortex of young rats. The FXR1 protein was detected in the perinuclear cytoplasm of cortical neurons (Figure 2A). The location of FXR1 precisely coincided with the signals of Thioflavin S (Figure 2A). Taking into consideration the fact that FXR1 forms SDS-resistant conformers and strongly colocalizes with Thioflavin S, we can conclude that this protein is present in rat brain in amyloid form under native conditions.

**Fig. 2.**
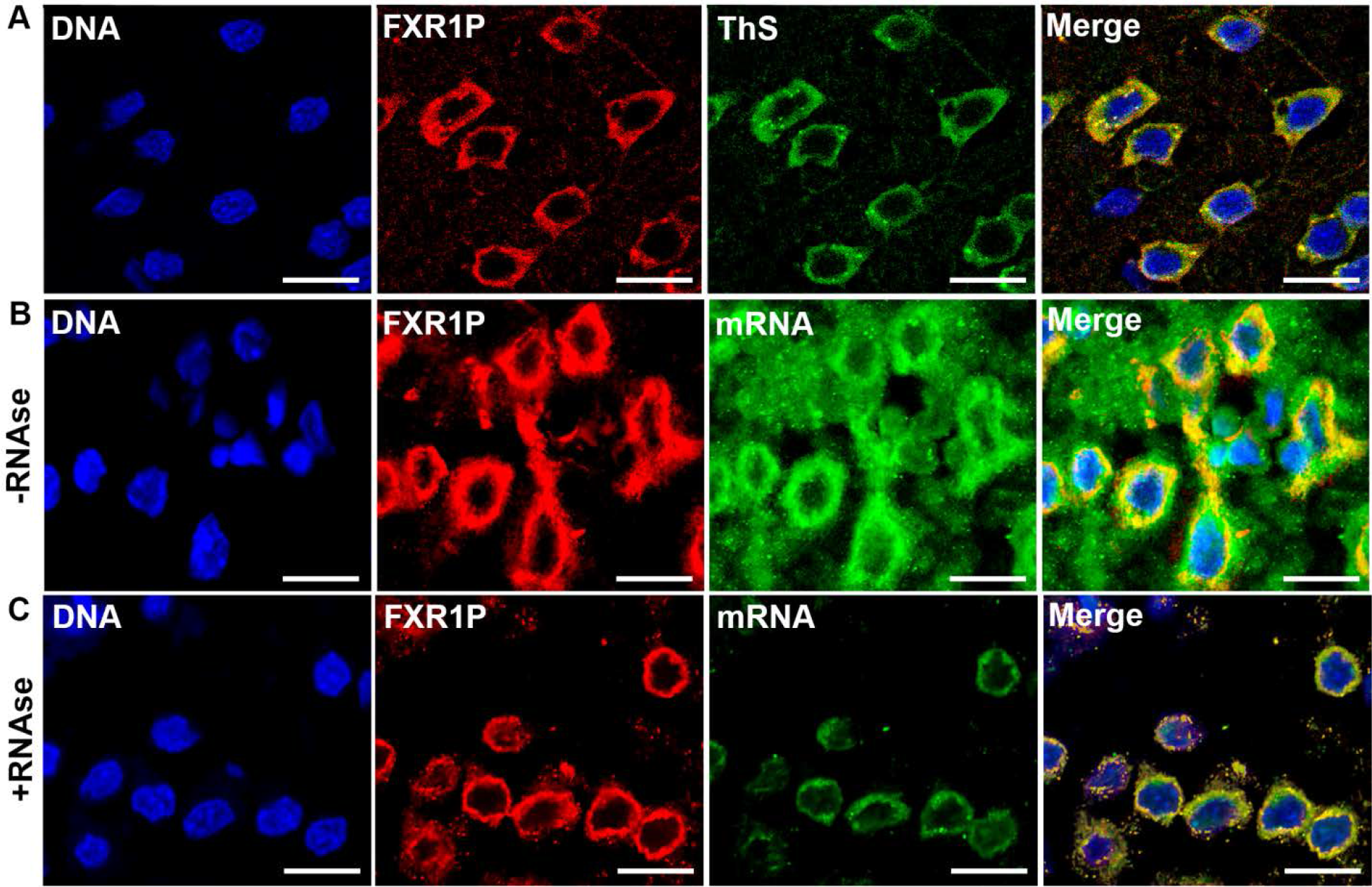
FXR1 colocalizes with Thioflavin S in cortical neurons, binds mRNA and protects mRNA from degradation. (A) FXR1 is present in the perinuclear cytoplasm of cortical neurons and colocalizes with amyloid-specific dye Thioflavin S. (B) FXR1 colocalizes with some portion of mRNA in the cytoplasm of cortical neurons. (C) mRNAs that are colocalized with FXR1 are detected after treatment with RNAse A, whereas other mRNAs degrade. Scale bar for sections A-C is 20 μm

It is known that FXR1 binds to different RNA molecules that form secondary structures and affects their level and translation efficiency (25). We suggested that amyloid conformers of FXR1 may protect RNA from action of ribonucleases that normally present in eukaryotic cells and take part in a gene expression control. To analyze the binding of FXR1 with mRNA, the brain cryosections were hybridized with biotinylated poly-dT, subsequently visualized by avidin-Alexa Fluor 488 and immunostained with anti-FXR1 antibody. As expected, we demonstrated that FXR1 colocalizes with a portion of mRNA in the cytoplasm of cortical neurons, whereas an essential part of mRNA is not associated with this protein (Figure 2B). Previously it was shown that mRNA molecules associated with FXR1 are resistant to RNase A treatment (26). Our finding that RNA is associated with amyloid conformers of FXR1 may explain this result. To analyze whether amyloid conformers of FXR1 affect mRNA stability, cryosections of the brain cortex were treated with RNase A in extremely high concentration (500 μg/ml). Then cryosections were hybridized with poly-dT and immunostained with anti FXR1-antibody. Molecules of mRNA that are colocalized with FXR1 were clearly detected after treatment with RNase A, whereas other mRNA molecules had completely degraded (Figure 2C). Thus, amyloid conformers of FXR1 bind RNA and prevent its degradation. Note, the members of ribonuclease A superfamily function in neurons (3).

### Aggregation of the FXR1 protein depends on its N-terminal region

Data obtained using the ArchCandy algorithm (17) show that the N-terminal part of FXR1 contains potentially amyloidogenic regions (Table 1). However, bioinformatic data are only predictive and require experimental verification. To identify the regions of FXR1 responsible for its aggregation, the sequences encoding FXR1(1-379 aa) and FXR1(380-568 aa) were PCR amplified from the total cDNA obtained from the brain-extracted mRNA of Wistar rats and fused in frame with the sequence encoding yellow fluorescent protein (YFP). The chimeric genes *FXR1(1-379)-YFP* and *FXR1(380-568)-YFP* were expressed under the control of the *CUP1* promoter in the *Saccharomyces cerevisiae* strain BY4742. As expected, FXR1(1-379)-YFP formed dot-like fluorescent foci in yeast cells (Figure 3A). The FXR1(380-568)-YFP protein exhibited diffuse fluorescence in the yeast cytoplasm (Figure 3A). Protein lysates were obtained from yeast cells expressing the chimeric proteins, centrifuged and separated into the soluble and insoluble fractions. The FXR1(1-379)-YFP protein was predominantly present in insoluble fraction, whereas the FXR1(380-568)-YFP was detected in soluble fraction (Figures 3B, 3C and 3D). Thus, the differential centrifugation analysis coupled with Western blot confirms the fluorescent microscopy data.

**Fig. 3.**
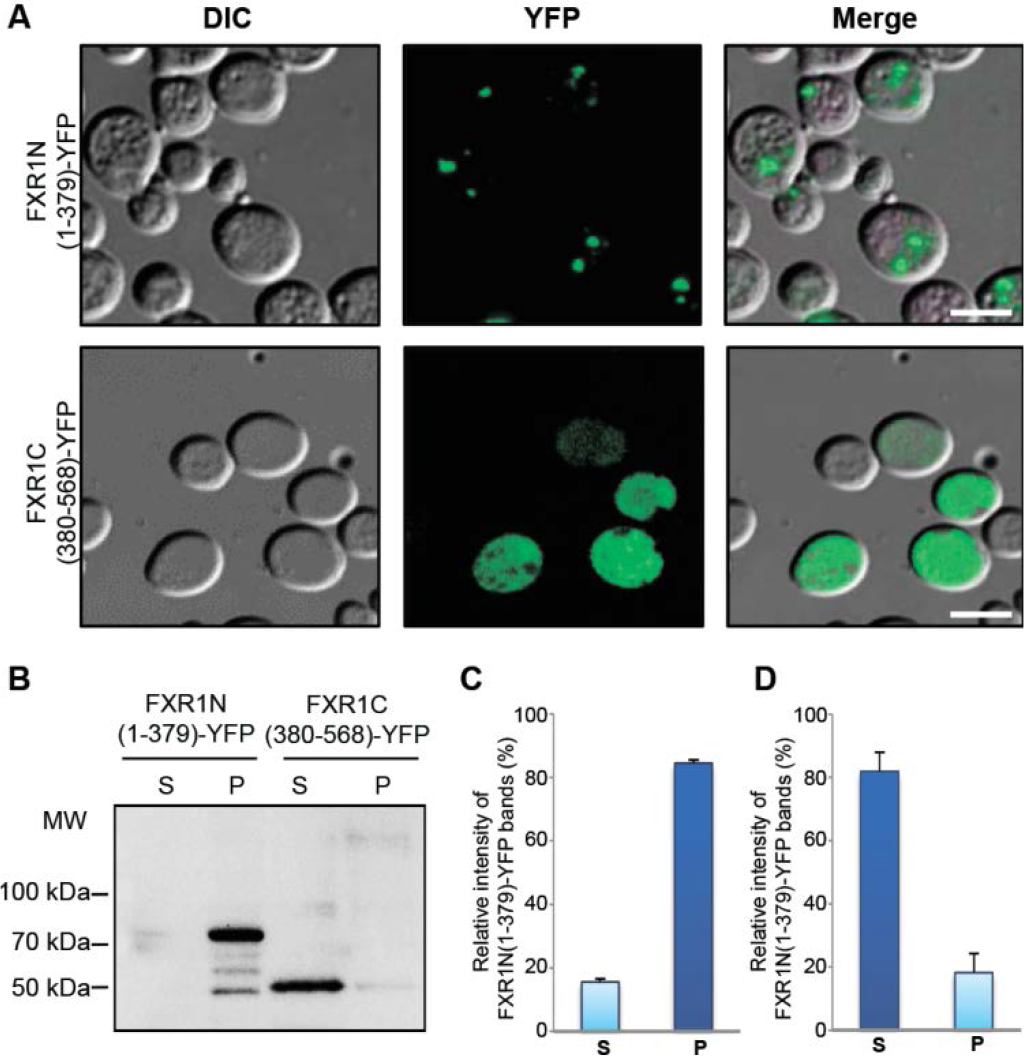
Aggregation of FXR1 protein depends on its N-terminal region. (A) FXR1N(1-379)-YFP protein forms visible aggregates, whereas FXR1C(380-568)-YFP is evenly distributed in cytoplasm. Scale bar, 10 μm. (B) The FXR1(1-379)-YFP protein expressed in yeast cells forms insoluble aggregates, whereas the FXR1(380-568)-YFP presents in soluble form. P – pellet fraction; S - supernatant fraction. (C) and (D) - Relative intensities of bands corresponding to monomers and aggregates of FXR1N(1-379)-YFP and FXR1C(380-568)-YFP represented as mean ± SEM, for three independent protein samples.

The amyloid properties of N-terminal domain of FXR1 were also analyzed *in vitro*. cDNA encoding the FXR1(1-379) fragment was synthetized from mRNA extracted from rat brain. This fragment was amplified and inserted into pET302 vector after 6xHis encoding sequence. The 6xHis-FXR1(1-379) recombinant protein immediately aggregated when placed into Tris buffer at 37°C, however microscopically discernible fibrils were detected only after 48 hours of incubation with slow rotation (Figure 4A). These fibrils bound Congo Red (Figure 4B) and manifested apple-green birefringence when assayed by polarization microscopy (Figure 4C).

**Fig. 4.**
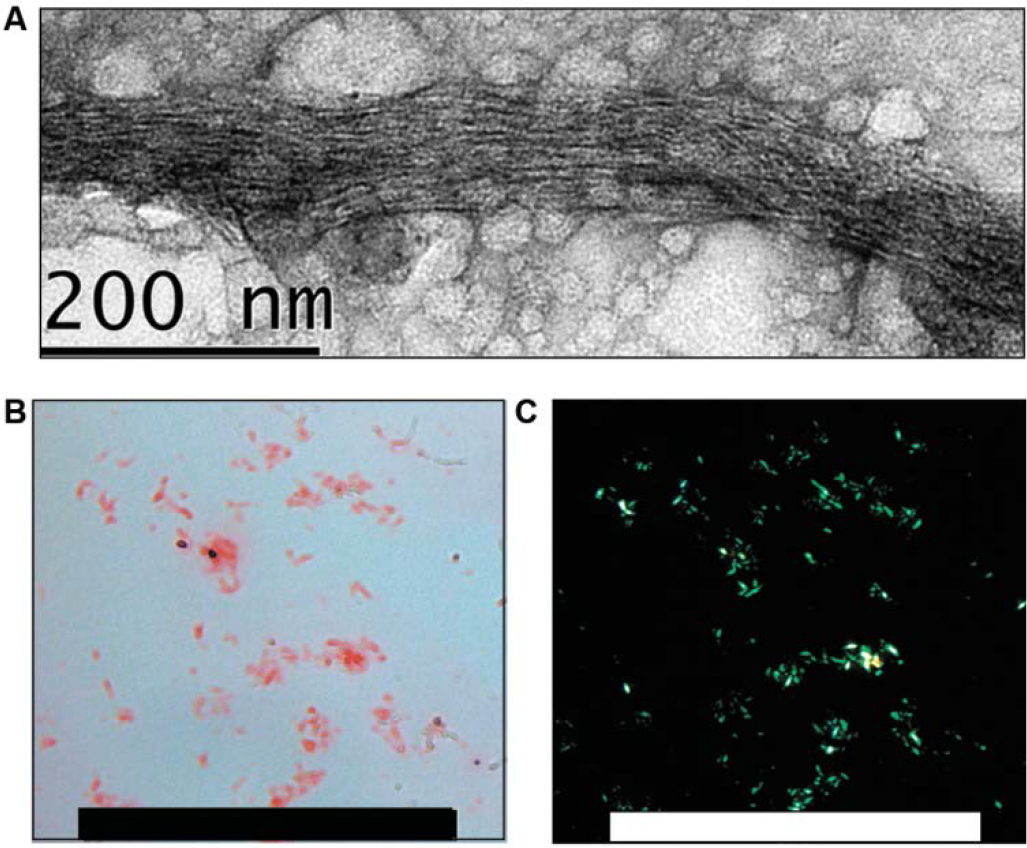
Recombinant protein 6xHis-FXR1(1-379) demonstrates amyloid properties in vitro. (A). Fibrils of 6xHis-FXR1(1-379) visualized by TEM. Scale bar, 200 nm. (B). Fibrils of 6xHis-FXR1(1-379) bind CR and (C) display apple-green birefringence when viewed between crossed polarizers. Scale bar, 100 μM.

To verify amyloid properties of N-terminal domain of FXR1, we also used the bacterial curli-dependent amyloid generator (C-DAG) system for production of extracellular amyloid fibrils (27, 28). This system relies on the ability of *E. coli* cells to generate surface associated amyloid fibrils (curli) composed of CsgA protein. This protein contains an N-terminal signal sequence (CsgA_SS_) that directs it to the cell surface and a C-terminal sequence that allows formation of extracellular fibrils (29, 30). Joining the signal sequence (CsgA_SS_) of the CsgA protein to heterologous amyloidogenic protein fragments directs their export to the cell surface where they can form fibrils. Since the C-DAG is applicable for the study of relatively short protein fragments (27), we analyzed the amyloidogenic properties of the N-terminal fragment of FXR1 spanning the first 337 amino acids. The *E. coli* VS39 strain was transformed with the pVS-FXR1(1-337) plasmid providing expression of CsgA_SS_ fused to the FXR1(1-337) fragment. The VS39 cells expressing CsgA_SS_-Sup35NM and CsgA_SS_-Sup35M proteins were used as positive and negative controls, respectively. We showed that expression of CsgA_SS_-FXR1(1-337) resulted in red colonies of transformants, although the red coloration was not as intense as in transformants expressing CsgA_SS_-Sup35NM (Figure S6A). The transformants expressing CsgA_SS_-Sup35M protein were pale on this medium (Figure S6A). CsgA_SS_-FXR1(1-337) and CsgA_SS_-Sup35NM proteins, but not CsgA_SS_-Sup35M, bound Congo Red and exhibited “apple-green” birefringence when examined between crossed polarizers (Figure S6B). This property is characteristic of amyloid fibrils (31). Using transmission electron microscopy we detected the extracellular fibrils of the CsgA_SS_-FXR1(1-337) and CsgA_SS_-Sup35NM proteins, but not of the CsgA_SS_-Sup35M protein (Figure S6C). Taking together, these data show that the N-terminal region of the FXR1 protein forms amyloid fibrils both *in vitro* and in the bacterial-based C-DAG system.

### N-terminal region of the FXR1 protein is evolutionary conserved

In order to analyze the evolutionary conservatism of FXR1, the ORF of corresponding gene was PCR amplified from the total cDNA obtained from brain-extracted mRNA of Wistar rats and sequenced (GenBank, accession number MG938503). The deduced amino acid sequence of the rat FXR1 protein was compared with canonical sequences of the FXR1 protein from different vertebrate species available in the UniProt database (http://www.uniprot.org). Since the amyloid properties of FXR1 depended on its N-terminal region, the comparative analysis was carried out separately for the N and C-terminal regions of this protein. The N-terminal sequence contains KH1 and KH2 RNA-binding motives, whereas the C-terminal region contains RGG (G quartet RNA motif) ^32^. The FXR1N(1-379 aa) sequences of rat and mouse are identical (Figure 5, Figure S7). They differ from human, macaque and cat sequences only by one amino acid – glutamic acid (E) in 270-th position is replaced by aspartic acid (D) (Figure S8). Both, E and D, belong to the class of the polar negatively charged amino acids, and this replacement is unlikely to change the amyloidogenic properties of the FXR1 protein. The N-terminal sequences of human, cat and macaque FXR1 proteins are identical. The FXR1N sequence of *Bos taurus* differs by two amino acids from the rat sequence (Figure 5, Figure S7). The N-terminal regions of FXR1 proteins of non-mammalian vertebrates differs from the rat FXR1N more significantly, but all of them contain two identically arranged potentially amyloidogenic regions (Figure 5). The C-terminal part of FXR1 of some species contains deletions before the RGG RNA-binding motive. No potentially amyloidogenic regions were revealed in FXR1C in all species analyzed (Figure S7). Thus, the N-terminal amyloid-forming region of FXR1 is highly conserved in the mammalian lineage and contains identically arranged amyloidogenic sequences in vertebrate species.

**Fig. 5.**
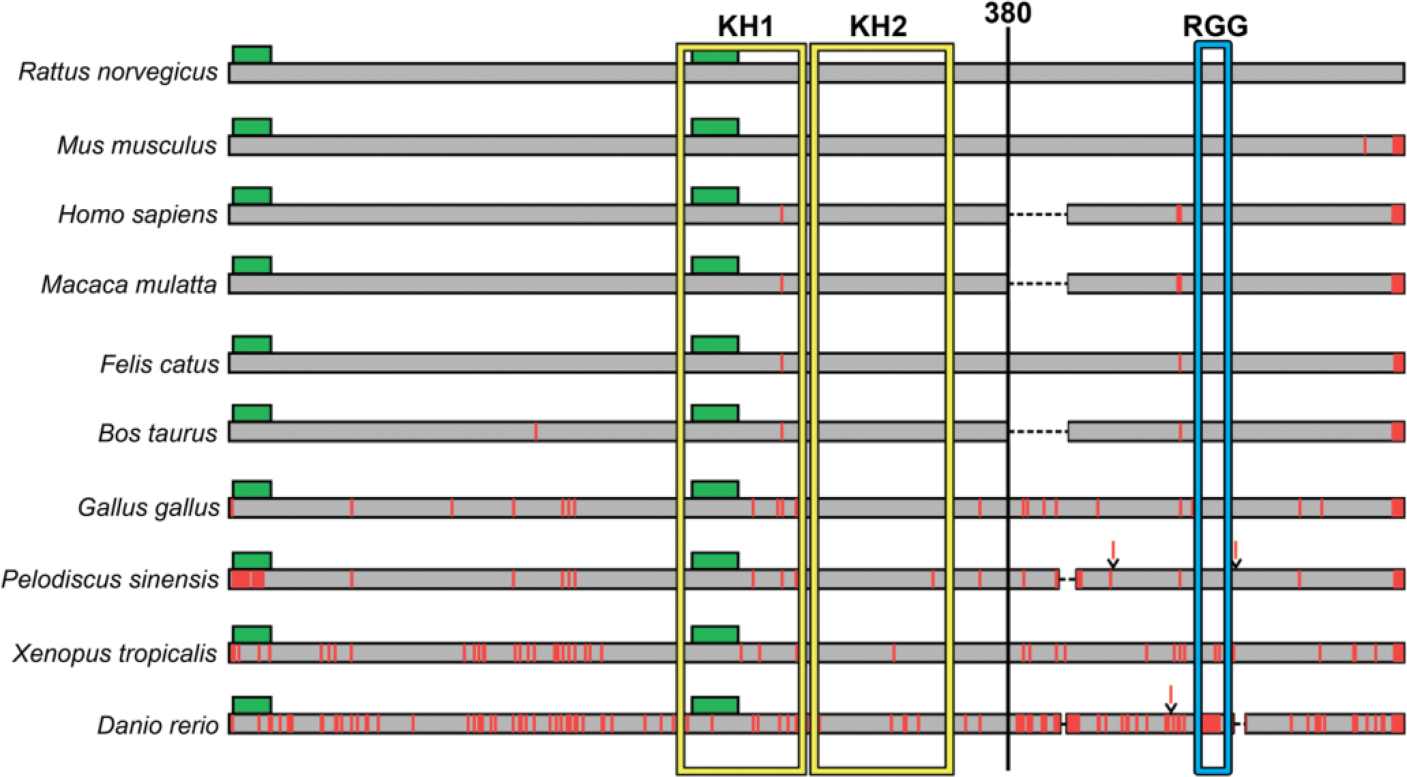
Diagram of the sequence alignment for the full-length FXR1 proteins from ten vertebrate species. The sequence of the FXR1 protein of Wistar rats is marked in gray. Potential amyloidogenic regions are marked as green rectangles. Positions of amino acids differing from those in the sequence of rat FXR1 are marked by a vertical red line. Insertions of amino acids are marked as red lines above the sequences. Deletions are indicated by a dashed black line. RNA binding domains KH1 and KH2 are enclosed in a yellow frame, and RGG domain – in a blue frame. See also Figure S7.

## Discussion

Proteomic screening, based on the universal biochemical properties of amyloids, provides us with new opportunities for the study of the prevalence and role of amyloids in nature. Using an original proteomic approach, we identified a set of proteins forming amyloid-like aggregates in the rat brain. The data obtained by PSIA-LC-MALDI approach opens up broad prospects for further identification of functional amyloids in mammalian brain. We focused on the analysis of amyloid properties of the FXR1 protein, but at the same time we identified several other promising candidates for the role of functional amyloids. The NSF, MBP, RIMS1 and STXB1 proteins were identified by mass-spectrometry in the SDS-resistant fraction of all analyzed brain samples (Table 1; Figure S1 – S5). All of them, except MBP, contain potentially amyloidogenic regions predicted by ArchCandy algorithm (Figure 1). The analysis of their amyloid properties will be the subject of our further research. Notably, there are some data suggesting that the myelin basic protein (MBP), that is a component of specialized membrane covering axons, may normally form amyloid structures in the brain (18). The CPEB3 protein was not identified in the SDS-resistant fraction in our proteomic screen. This is not surprising, since it was shown in the mouse model that CPEB3 aggregates only in the brains of specifically stimulated animals (11).

Here we provided comprehensive evidences that the FXR1 protein forms amyloid conformers *in vivo* in rat brain under native conditions. Moreover, amyloid conformers of FXR1 protect RNA from degradation in cortical neurons. Previously it was shown that FXR1 binds and stabilizes certain RNAs, such as mRNAs encoding desmosomal proteins and non-coding telomerase RNA (26, 33). The molecular basis of this phenomenon remained unknown. As we demonstrated, FXR1 forms amyloid conformers that bind RNA molecules and protect them from the RNase-A mediated degradation in the cytoplasm of cortical neurons (Figure 4). Important to note that the members of ribonuclease A superfamily function in various cells, including neurons (3).

We showed that amyloid conformers of FXR1 are present in both soluble (oligomeric) and insoluble (pellet) fractions in rat brain (Figure 2A and 2B). There are evidences that FXR1 in soluble and insoluble states differently affects the translation of mRNA. Previously it was shown that FXR1 forms in cell culture small soluble RNP particles that induce cell-cycle arrest in serum-starvation conditions. These particles activate translation of mRNAs containing AU-rich signal sequence (34). In serum-grown cells, FXR1 appears in large insoluble cytoplasmic bodies that are associated with translation silencing (35, 36). Probably, protection of untranslated mRNA in cytoplasmic bodies is important for the regulation of the cell response to different exposures. Thus, amyloid oligomers and insoluble aggregates of FXR1 both protect mRNA from degradation, but they differently regulate mRNA expression.

Recently it was shown that the level of FXR1 production in cortical neurons affects long-term memory, mood and emotions. Removal of FXR1 from the forebrain of postnatal mice selectively enhances long-term memory (1), whereas overexpression of FXR1 in the prefrontal cortex leads to an antidepressant-like effect and has a positive effect on emotional stability (2). The overexpression of FXR1 and mood stabilization can be induced by treatment with lithium, which is a prototypal psychotropic drug (2). It is possible that some treatments or stress inducing factors can provoke not only alterations of FXR1 expression, but also its transition between an oligomeric state and insoluble aggregates. Such transitions may represent the mechanism for regulation of protein synthesis. We are planning to evaluate the effects of psychotropic impacts on FXR1 aggregation and mRNA translation in future research.

Bioinformatic algorithm predicted that the N-terminal part of FXR1 contains potentially amyloidogenic regions (Table 1). The amyloid properties of N-terminal fragment of FXR1 were confirmed in yeast-based system, *in vitro* and in the bacterial-based C-DAG system (Figure 4). Moreover, N-terminal fragment of FXR1 is highly conserved across mammals and contains identically arranged potentially amyloidogenic sequences in evolutionary distant vertebrates (Figure 5, Figure S7). These data suggest that the ability of amyloid conformers of FXR1 to protect specific mRNAs is evolutionary conserved, especially in the mammalian lineage including human. Finally, we note that identification of functional amyloids in mammalian brain requires a review of strategy for amyloidosis treatment. The strategy aimed at the development of universal anti-amyloid drugs is unpromising. Such potential drugs should prevent or suppress formation of pathological aggregates of a certain protein, but not affect functional amyloids that regulate cognitive processes.

## Materials and Methods

### Animals

Six-month-old male Wistar rats were purchased in Rappolovo breeding colony (Saint-Petersburg, Russia). All animal experiments were conducted in accordance with the Treaty of Lisbon amending the Treaty on European Union and the Treaty establishing the European Community and entered into force on 1 December 2009. Experiments were approved by the Ethical Committee for Animal Research of St. Petersburg State University (conclusion # 131-03-6). Euthanasia of animals was carried out immediately after delivery to Saint-Petersburg State University.

### Proteomic screening and identification of proteins forming amyloid‑like aggregates

Proteomic screening and identification of proteins forming amyloid‑like aggregates in brain of *Rattus norvegicus* was performed using PSIA–LC–MALDI approach described recently (12) with modification (more details are provided in *SI Appendix*).

### Rat brain slices and homogenization

For immunohistochemistry and fluorescent in situ hybridization rat brains were extracted, washed with PBS and fixed in 4% PFA for 3 hours. After fixation brains were embedded in FSC22 compound (Leica), frozen in liquid nitrogen and stored at −70°C. Brains were sectioned to 20 μm thick slices on a freezing microtome CM1850UV (Leica). For PSIA–LC–MALDI analysis brains after extraction were immediately frozen in liquid nitrogen and homogenized using a cryogenic laboratory mill Freezer/Mill 6870 (SPEX SamplePrep).

### <u>I</u>mmunohistochemistry, fluorescent in situ hybridization and confocal microscopy

Immunohistochemistry and histological staining was performed on rat brain sections. Enzymatic treatment was performed with RNase A (500μg/ml, Thermo Fisher Scientific, USA) for one hour at 37° degrees (details are provided in *SI Appendix*).

### Cloning of the *FXR1* gene fragments

Total rat RNA was extracted from brain homogenates using TRIzol reagent (Invitrogen) according to the manufacturer protocol. cDNA synthesis was performed with SuperScript III Reverse Transcriptase (Invitrogen). cDNA was further used for *FXR1* fragments synthesis. All primers are listed in Table S2 (see also *SI Appendix*).

### Protein Expression and Purification

FXR1(1-379) was expressed as His- tagged fusion protein in Rosetta (DE3) pLysS cells (Novagen) in LB medium. Cells were lysed in 0.05 M Tris (pH 7.6), 0.15 M NaCl and complete protease inhibitor cocktail (Roche) by ultrasonication. Inclusion bodies were dissolved in 8M urea, 20mM Na phosphate, 0,25M NaCl, 5 mM imidazole and 1mM βME. The cleared lysate was loaded onto a gravity NiNTA column (Qiagen), washed with 8M urea, 20mM Na phosphate, 0,25M NaCl, 5 mM imidazole and 1mM β-mercaptoethanol, and eluted in 8M urea, 20mM Na phosphate, 0,5M NaCl, 500 mM imidazole and 1mM βME. The purity of the recombinant protein was verified by SDS-PAGE, protein concentration was measured using Qubit 2 Fluorometer (Thermo Scientific).

### *In Vitro* Fibril Formation Assay

The experiment was performed as described previously (37). In brief, purified FXR1(1-379) protein solution diluted to 5 μM in a buffer containing 20 mM Tris (pH 8.0), 1 mM βME was incubated at 37°C under agitation for 48 hr with slow rotation. Fibril formation was verified by TEM and Congo Red staining.

### Electron microscopy

TEM images were recorded on a Jeol JEM-2100 microscope. Negatively-stained samples were prepared on formvar film 300 mesh copper grids (Electron Microscopy Sciences). A 10 μl aliquot of fibril solution (from in vitro experiment) or bacterial culture solution (from C-DAG experiment) were adsorbed to the formvar film for one minute, blotted, washed twice with 10 μl of water for 10 s, stained with 10 μl of 1% uranyl acetate for 1 minute and dried in air.

### Congo Red staining of fibrils

10μL of the aggregated protein solution was put onto a glass microscope slide, air-dried, stained with 50μL of Congo Red water solution (2,5 mg/ml), washed with water and analyzed between cross polarizers on the inverted microscope Leica DMI6000 B fitted with cross polarizers. Images were acquired using the Leica Application Suite software.

### Protein analysis

Preparation of protein lysates was performed as described previously (38) (more details are provided in *SI Appendix*).

### Yeast strains and growth conditions

The *S. cerevsiae* strain BY4742 (*36*) [*MATα his3Δ1 leu2Δ0 lys2Δ0 ura3Δ0*] was used for analysis of the FXR1 fragments aggregation. Standard yeast genetic techniques, media, and cultivation conditions were used. Yeast cultures were grown at 30°C. 150 μM copper sulfate (CuSO_4_) was added to synthetic medium to induce the expression of genes under the *P*_*CUP1*_ promoter.

### Analysis of amyloid fibril formation in the bacteria‑based system C‑DAG

The analysis of amyloid fibril formation of FXR1(1-337aa) fragment in the bacteria‑based C‑DAG system was performed as described earlier (27, 28). *E. coli* strain VS39 was transformed with the plasmid pVS-FXR1(1-337) coding for FXR1(1-337) protein fused to CsgA signal sequence. VS39 transformants with pVS72 and pVS105 encoding the CsgA_SS_-Sup35NM and CsgASS-Sup35M proteins were used as positive and negative controls of amyloid generation, respectively. Tests for colony color phenotype and CR birefringence were performed as described in (27). To perform CR birefringence analysis, suspensions of the cells grown on CR-containing medium were spotted onto a glass slide and analyzed between cross polarizers on the inverted microscope Leica DMI6000 B fitted with cross polarizers. Images were acquired using the Leica Application Suite software. To perform transmission electron microscopy analysis, spots of the transformants were grown for 5 days at 22°C on the inducing medium without Congo Red (LB supplemented with 100 mg/l ampicillin, 25 mg/l chloramphenicol, 0.2% w/v L- arabinose and 1 mM IPTG).

## Acknowledgements

The authors acknowledge Dr. A.I. Aleksandrov for critical reading of the manuscript. Special thanks to Dr. A. Hochschild for providing the bacterial C-DAG system. We are grateful to Dr. Vol’nova A.B., and Dr. V.S. Ioffe, Dr. S.A.Galkina, Dr. G.O.Cherepanov, A.N. Lyholay and M.G. Vorobiev for technical support and helpful discussion. The authors acknowledge St. Petersburg State University for opportunity to use facilities of the Research Resource Center for Molecular and Cell Technologies and the Resource Centers “CHROMAS”. This work was supported by state program # 0112-2016-0015 to A.P.G., by the grants of SPbSU to A.P.G. and S.G.I. and grant of SpbSU to Laboratory of Amyloid Biology, proteomic screening partially was supported by RFBR according to the research project № 18-34-00419.

