## Supplementary materials for "Amyloid conformers of the FXR1 protein prevent mRNA degradation in cortical neurons"

Corresponding author: Alexey P. Galkin

This PDF file includes:

Supplementary Materials and Methods

Figs S1 to S7

Tables S1 to S2

References for SI reference citations

### Supporting Methods

#### Proteomic screening and identification of proteins forming amyloid-like aggregates

Schematic diagram of the methodology is shown in the Figure S8. Brain homogenates were solubilized in Tris-buffered saline (TBS) (30 mM Tris-HCl, pH 7.4, 150 mM NaCl), supplemented with 10 mM PMSF and Complete Protease Inhibitor (Roche). The obtained lysates were precleared (2500 g, 10 min, 4°C) and then fractionated by ultracentrifugation (151000 g, 2 h, 4°C). Pellets containing protein aggregates were resuspended in TBS with 200 µg/ml RNAse A, incubated for 15 min and treated with 1% SDS for 8 hours at room temperature. Then, detergent-resistant protein complexes were separated from monomeric proteins by ultracentrifugation at 151000 g, 8 h, 18°C through 25% sucrose-TBS cushion with 0.1% SDS. Pellets were suspended in water, sedimentated again for at 151000 g (2 h, 4°C) and boiled in SDS-PAGE loading buffer. Then, detergents and salts were removed from the samples using HiPPR Detergent Removal columns (Thermo Scientific, USA) and Zeba Desalting columns (Thermo Scientific, USA), respectively, according to the manufacturers' protocols. Samples (volume 50 µl, total protein concentration 0.2–0.4 mg/ml) were supplemented with 1 µl of freshly prepared 50 mM DTT in 50 mM ammonium bicarbonate, incubated for 15 min at 50°C, supplemented with 1 µl 100 mM iodoacetamide in 50 mM ammonium bicarbonate and incubated for 15 min at 20°C in the dark. Then the samples were supplemented with 1 µl DTT to inactivate iodoacetamide and 5 µl trypsin (10 ng/µl; Sigma) and incubated overnight at 37°C. The trypsin was inactivated by adding 0.5 µl 10% TFA followed by centrifuging for 30 min (20,000g, 4°C). The final peptide mixtures were loaded (1 µl) onto an Acclaim PepMap 300 HPLC reverse-phase column (150 mm, 75 µm, particle size 5 µm; Thermo Scientific, USA) and separated in an acetonitrile gradient (2–90%) during 45 min using an UltiMate 3000 UHPLC RSLC nano high-performance nanoflow liquid chromatograph (Dionex, USA). Peptide fractions were collected every 10 s and loaded onto a 384-sample MTP AnchorChip 800/384 microtiter plate (Bruker Daltonics) using a Proteineer fc II spotter (Bruker Daltonics).

Peptides were identified using the Ultraflextreme MALDI-TOF/TOF mass spectrometer (Bruker Daltonics, DE). MS-spectra for each peptide fraction were determined and analyzed using WARP-LC software. An array of unique peptides characterized by specific retention time, charge, and molecular weight was determined. MS/MS-analysis was performed for these peptides in fractions (spots) with maximal concentration (peak intensity) of these peptides. Match between the experimental spectra and corresponding proteins was analyzed using Mascot version 2.4.2 software (Matrix Science; http://www.matrixscience.com) in the UniProt database (http://www.uniprot.org/) restricted to *Rattus norvegicus*. α-cyano-4-hydroxycinnamic acid was used as a matrix. During analysis, preset parameters of "Mass tolerance" were used (precursor mass tolerance 100 ppm, fragment mass tolerance 0.9 Da). As a standard sample, Peptide Calibration Standard II 8222570 (Bruker Daltonics) was applied. Carboxymethylation of cysteine, partial oxidation of methionine, and one skipped trypsinolysis site were considered as permissible modifications. The BioTools software (Bruker, Bremen, Germany) was used for manual validation of protein identification.

**Immunohistochemistry, fluorescent in situ hybridization and confocal microscopy**

Histological staining for amyloids was done using 1% Thioflavin S (ThioS, Sigma, USA) solution. The primary Anti-FXR1 antibody ab129089 (1:200, Abcam, Cambridge, MA) and secondary antibody Goat anti-Rabbit IgG (H+L) conjugated with Alexa Fluor 546 (1:500, Thermo Fisher Scientific, USA) were used. PolyA mRNA was detected with biotinylated oligo(dT)20 (Beagle, Russia) followed by avidin conjugated with Alexa Fluor 488 (1: 200, Thermo Fisher Scientific, USA). The slides were also subjected to DAPI nuclear staining. Microscopic analysis was performed using a TCS SP5 confocal microscope (Leica Microsystems, Germany) and "Leica Application Suite X 3.3.0.16799" software.

**Cloning of the *FXR1* gene fragments**

The fragment of *FXR1* gene coding for 1-379 aa for expression in *E. coli* was amplified with *fxr1 EcoRI forward* and *fxr1(379) BamHI reverse* primers and this fragment was inserted into pET302 vector to obtain the pET302-FXR1(1-379) plasmid. The fragment of *FXR1* gene coding for 1-379 aa for expression in *S. cerevisiae* as amplified with *fxr1 HindIII forward* and *fxr1 (YFP)BamHI reverse* primers. The *fxr1 (380) HindIII forward* and *fxr1 (568) BamHI reverse* primers were used for amplification of the fragment of *FXR1* gene coding for 380-568 aa. Both fragments were inserted in pCUP-YFP vector to obtain pCUP-FXR1(1-379)-YFP and pCUP-FXR1(380-568)-YFP plasmids. The fragment of the *FXR1* gene coding for 1-337 aa for expression in C-DAG system was amplified with *fxr1 NotI forward* and *fxr1 XbaI reverse* primers and this fragment was inserted into pVS72 vector to obtain the pVS-FXR1(1-337) plasmid.

**Protein analysis**

Rat brain lysates were fractionated for 25 min at 9800 g, +4ºC and the insoluble fraction was collected. The supernatant was loaded on the Amicon Ultra 100K filter (Merck, Millipore) and further divided into two fractions: less than 100 kDa and more than 100 kDa. The proteins from three selected fractions were analysed by Western blotting. The Fxr1 protein was detected with the primary Anti-FXR1 antibody ab129089 (Abcam) and secondary Goat Anti-Rabbit IgG H&L(ab205718) (HRP) (Abcam). Chemiluminescent detection was performed using the Amersham ECL Prime Western Blotting Detection Reagent (GE Healthcare, USA). Relative intensity of bands corresponding to FXR1 monomers and SDS-resistant conformers is represented as mean ± SEM for three independent brain samples.Fractionation of yeast cell lysates was performed according to (1). Cell lysate was centrifuged for 25 min at 9800 g, +4ºC, then supernatant was transferred to a new tube and debris was suspended in equal buffer volume. Both supernatant and debris were analysed by Western blotting. Semi-denaturing detergent agarose gel electrophoresis (SDD-AGE) (2, 3) was performed using 1% agarose gel. Before loading onto a gel, protein extracts were treated for 10 min with 1% SDS at room temperature. Then, the extract was subjected to SDD-AGE and transferred onto Immobilon-P PVDF membrane (GE Healthcare, USA). Proteins fused with YFP were detected with polyclonal chicken primary antibodies against GFP (ab13970) (Abcam, Great Britain) and secondary Goat Anti-Chicken IgY H&L (HRP) (ab6877) (Abcam). Chemiluminescent detection was performed using the Amersham ECL Prime Western Blotting Detection Reagent (GE Healthcare, USA). Relative intensities of bands corresponding to monomers and aggregates of FXR1N(1-379)- YFP and FXR1C(380-568)-YFP was represented as mean ± SEM for three independent protein samples.

### Supporting Tables, Figures, and Legends

NSF  Vesicle-fusing ATPase OS=Rattus norvegicus GN=Nsf PE=1 SV=1 NSF_RAT MH+(mono):1.008  MH+ (avg): 1.008

Tolerance (Da):0.500 Number of Peaks:775

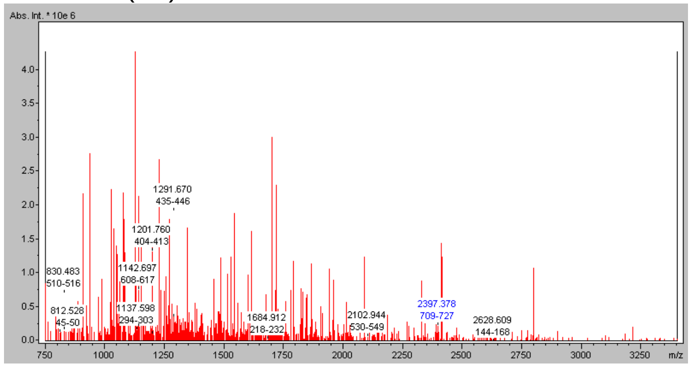


Tolerance (Da)  0.900   Number of Peaks: 1726

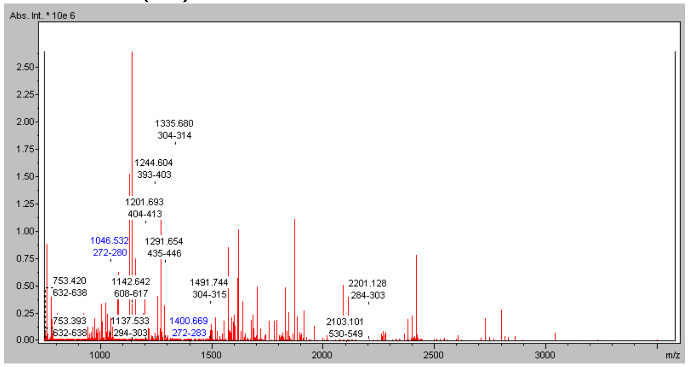


Tolerance (Da):0.500 Number of Peaks: 616

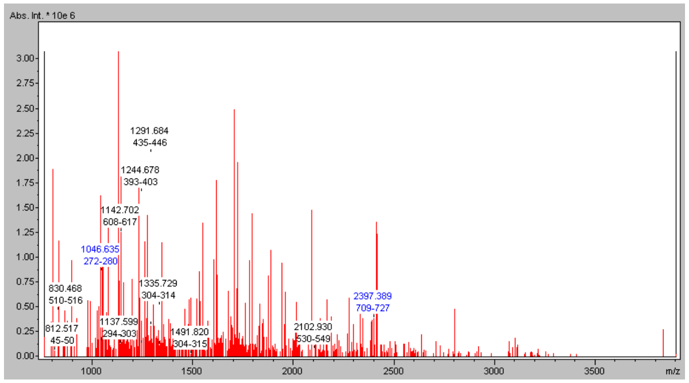


1 10 20 30 40 50 60
MAGRTMQAAR CPTDELSLSN CAVVNEKDYQ SGQHVMVRTS PNHKYIFTLR THPSVVPGCI
70 80 90 100 110 120
AFSLPQRKWA GLSIGQDIEV ALYSFDKAKQ CIGTMTIEID FLQKKNIDSN PYDTDKMAAE
130 140 150 160 170 180
FIQQFNHQAF SVGQQLVFSF NDKLFGLLVK DIEAMDPSIL KGEPASGKRQ KIEVGLVVGN
190 200 210 220 230 240
SQVAFEKAEN SSLNLIGKAK TKENRQSIIN PDWNFEKMGI GGLDKEFSDI FRRAFASRVF
250 260 270 280 290 300
PPEIVEQMGC KHVKGILLYG PPGCGKTLLA RQIGKMLNAR EPKVVNGPEI LNKYVGESEA
310 320 330 340 350 360
NIRKLFADAE EEQRRLGANS GLHIIIFDEI DAICKQRGSM AGSTGVHDTV VNQLLSKIDG
370 380 390 400 410 420
VEQLNNILVI GMTNRPDLID EALLRPGRLE VKMEIGLPDE KGRLQILHIH TARMRGHQLL
430 440 450 460 470 480
SADVDIKELA VETKNFSGAE LEGLVRAAQS TAMNRHIKAS TKVEVDMEKA ESLQVTRGDF
490 500 510 520 530 540 550 560 570 580 590
LASLENDIKP AFGTNQEDYA SYIMNGIIKW GDPVTRVLDD GELLVQQTKN SDRTPLVSVL LEGPPHSGKT ALAAKIAEES NFPFIKICSP DKMIGFSETA KCQAMKKIFD
600 610 620 630 640 650 660 670 680 690 700
DAYKSQLSCV VVDDIERLLD YVPIGPRFSN LVLQALLVLL KKAPPQGRKL LIIGTTSRKD VLQEMEMLNA FSTTIHVPNI ATGEQLLEAL ELLGNFKDKE RTTIAQQVKG
710 720 730 740
KKVWIGIKKL LMLIEMSLQM DPEYRVRKFL ALMREEGASP LDFD

**Fig. S1. Mass spectrometry identification data of the NSF protein (see also Table S1) in three brain samples of six-month-old male rats.** The amino acid sequence of the NSF protein is shown. The peptides identified by mass spectrometry are indicated in yellow.

MBP Myelin basic protein OS=Rattus norvegicus GN=Mbp PE=1 SV=3 MBP_RAT MH+ (mono):1.008 MH+ (avg): 1.008

Tolerance (Da):0.500 Number of Peaks:775

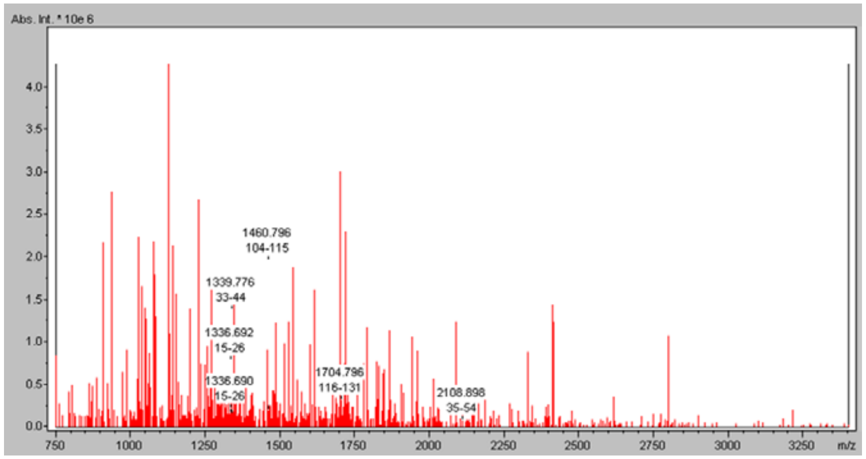


Tolerance (Da) 0.900 Number of Peaks: 1726

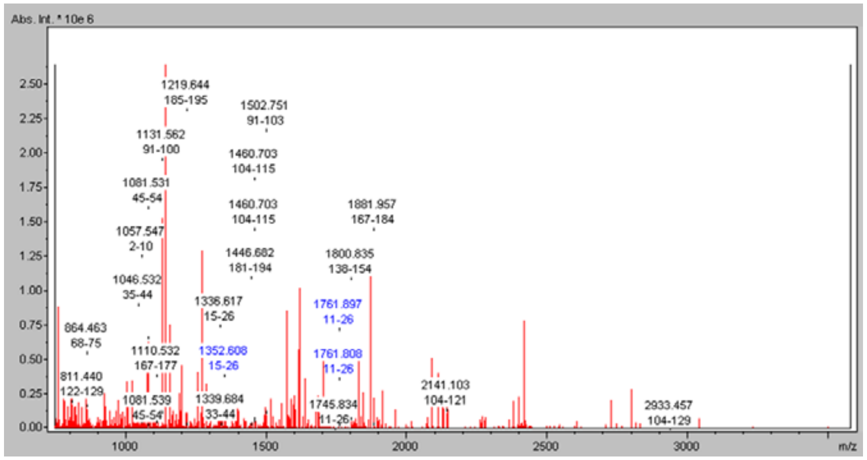


Tolerance (Da):0.500 Number of Peaks: 616

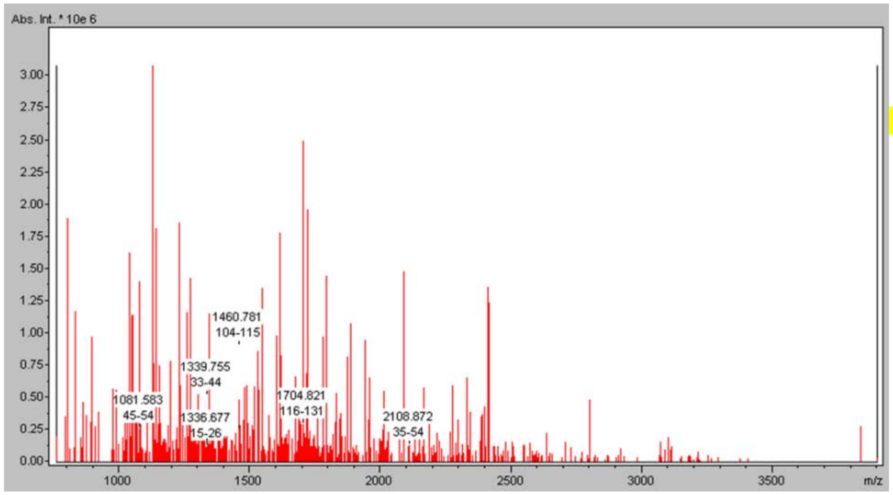


1 10 20 30 40 50
MASQKRPSQR HGSKYLATAS TMDHARHGFL PRHRDTGILD SIGRFFSGDR
60 70 80 90 100
GAPKRGSGKV PWLKQSRSPL PSHARSRPGL CHMYKDSHTR TTHYGSLPQK
110 120 130 140 150
SQRTQDENPV VHFFKNIVTP RTPPPSQGKG RGLSLSRFSW GAEGQKPGFG
160 170 180 190
YGGRASDYKS AHKGFKGAYD AQGTLSKIFK LGGRDSRSGS PMARR

**Fig. S2. Mass spectrometry identification data of the MBP protein (see also TableS1) in three brain samples of six-month-old male rats.** The amino acid sequence of the MBP protein is shown. The peptides identified by mass spectrometry are indicated in yellow.

RIMS1 Regulating synaptic membrane exocytosis protein 1 OS=*Rattus norvegicus* GN=Rims1 PE=1 SV=1 RIMS1_RAT

MH+ (mono):1.008 MH+ (avg): 1.008

Tolerance (Da):0.500 Number of Peaks:775

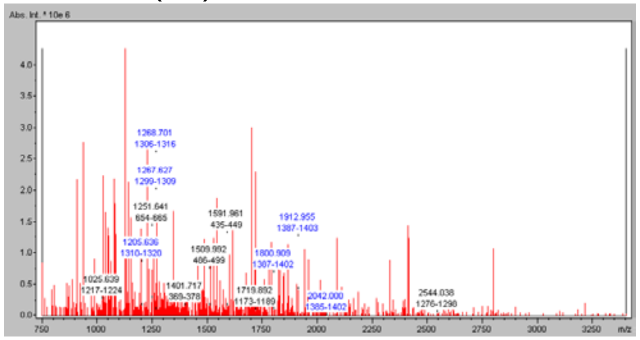

Tolerance (Da) 0.900 Number of Peaks: 1726

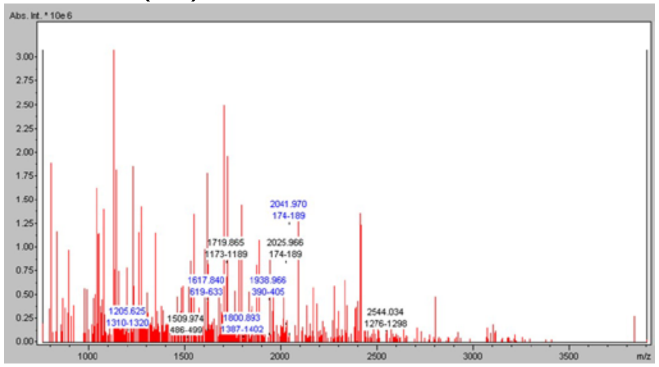

Tolerance (Da):0.500 Number of Peaks: 616

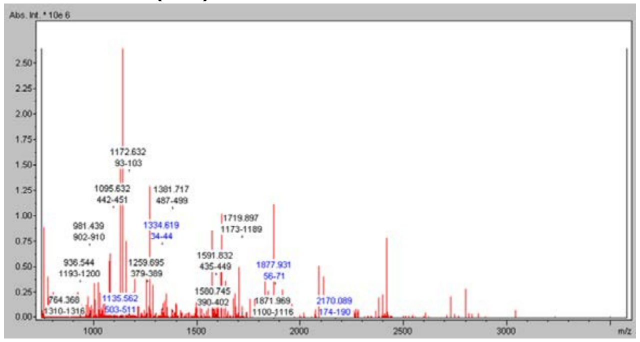

1 10 20 30 40 50 60 70 80
MSSAVGPRGP RPPTVPPPMQ ELPDLSHLTE EERNIIMAVM DRQKEEEEKE EAMLKCVVRD MAKPAACKTP RNAESQPHQP
90 100 110 120 130 140 150 160
PLNIFRCVCV PRKPSSEEGG PERDWRLHQQ FESYKEQVRK IGEEARRYQG EHKDDAPTCG ICHKTKFADG CGHLCSYCRT
170 180 190 200 210 220 230 240
KFCARCGGRV SLRSNNEDKV VMWVCNLCRK QQEILTKSGA WFFGSGPQQP SQDGTLSDTA TGAGSEVPRE KKARLQERSR
250 260 270 280 290 300 310 320
SQTPLSTAAV SSQDTATPGA PLHRNKGAEP SQQALGPEQK QASRSRSEPP RERKKAPGLS EQNGKGGQKS ERKRVPKSVV
330 340 350 360 370 380 390 400
QPGEGIADER ERKERRETRR LEKGRSQDYS DRPEKRDNGR VAEDQKQRKE EEYQTRYRSD PNLARYPVKA PPEEQQMRMH
410 420 430 440 450 460 470 480
ARVSRARHER RHSDVALPHT EAAAAAPAEA TAGKRAPATA RVSPPESPRA RAAAAQPPTE HGPPPPRPAP GPAEPPEPRV
490 500 510 520 530 540 550 560
PEPLRKQGRL DPGSAVLLRK AKREKAESML RNDSLSSDQS ESVRPSPPKP HRPKRGGKRR QMSVSSSEEE GVSTPEYTSC
570 580 590 600 610 620 630 640
EDVELESESV SEKGDLDYYW LDPATWHSRE TSPISSHPVT WQPSKEGDRL IGRVILNKRT TMPKESGALL GLKVVGGKMT
650 660 670 680 690 700 710 720 730 740 750 760 770 780
DLGRLGAFIT KVKKGSLADV VGHLRAGDEV LEWNGKPLPG ATNEEVYNII LESKSEPQVE IIVSRPIGDI PRIPESSHPP LESSSSSFES QKMERPSISV ISPTSPGALK DAPQVLPGQL SVKLWYDKVG HQLIVNVLQA
790 800 810 820 830 840 850 860 870 880 890 900 910 920
TDLPPRVDGR PRNPYVKMYF LPDRSDKSKR RTKTVKKLLE PKWNQTFVYS HVHRRDFRER MLEITVWDQP RVQDEESEFL GEILIELETA LLDDEPHWYK LQTHDESSLP LPQPSPFMPR RHIHGESSSK KLQRSQRISD
930 940 950 960 970 980 990 1000 1010 1020 1030 1040 1050 1060
SDISDYEVDD GIGVVPPVGY RASARESKAT TLTVPEQQRT THHRSRSVSP HRGDDQGRPR SRLPNVPLQR SLDEIHPTRR SRSPTRHHDA SRSPADHRSR HVESQYSSEP DSELLMLPRA KRGRSAESLH MTSELQPSLD
1070 1080 1090 1100 1110 1120 1130 1140 1150 1160 1170 1180 1190 1200
RARSASTNCL RPDTSLHSPE RERHSRKSER CSIQKQSRKG TASDADRVLP PCLSRRGYAT PRATDQPVVR GKYPTRSRSS EHSSVRTLCS MHHLAPGGSA PPSPLLLTRT HRQGSPTQSP PADTSFGSRR GRQLPQVPVR
1210 1220 1230 1240 1250 1260 1270 1280 1290 1300 1310 1320 1330 1340
SGSIEQASLV VEERTRQMKV KVHRFKQTTG SGSSQELDHE QYSKYNIHKD QYRSCDNASA KSSDSDVSDV SAISRASSTS RLSSTSFMSE QSERPRGRIS SFTPKMQGRR MGTSGRAIIK STSVSGEIYT LERNDGSQSD
1350 1360 1370 1380 1390 1400 1410 1420 1430 1440 1450 1460 1470 1480
TAVGTVGAGG KKRRSSLSAK VVAIVSRRSR STSQLSQTES GHKKLKSTIQ RSTETGMAAE MRKMVRQPSR ESTDGSINSY SSEGNLIFPG VRVGPDSQFS DFLDGLGPAQ LVGRQTLATP AMGDIQIGME DKKGQLEVEV
1490 1500 1510 1520 1530 1540 1550 1560 1570 1580 1590 1600 1610
IRARSLTQKP GSKSTPAPYV KVYLLENGAC IAKKKTRIAR KTLDPLYQQS LVFDESPQGK VLQVIVWGDY GRMDHKCFMG VAQILLEELD LSSMVIGWYK LFPPSSLVDP TLAPLTRRAS QSSLESSSGP PCIRS

**Fig. S3. Mass spectrometry identification data of the RIMS1 protein (see also Table S1) in three brain samples of six-month-old male rats.** The amino acid sequence of the RIMS1 protein is shown. The peptides identified by mass spectrometry are indicated in yellow.

STXBP1 Syntaxin-binding protein 1 OS=Rattus norvegicus GN=Stxbp1 PE=1 SV=1 STXB1_RAT MH+ (mono):1.008 MH+ (avg): 1.008

Tolerance (Da):0.500 Number of Peaks:775

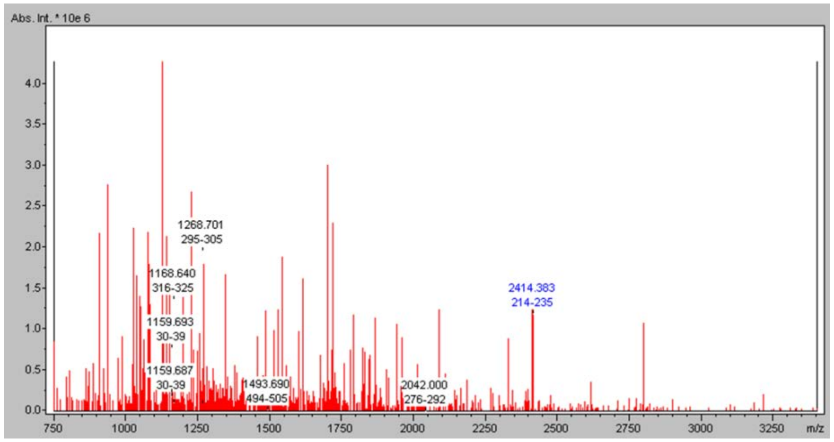


Tolerance (Da) 0.900 Number of Peaks: 1726

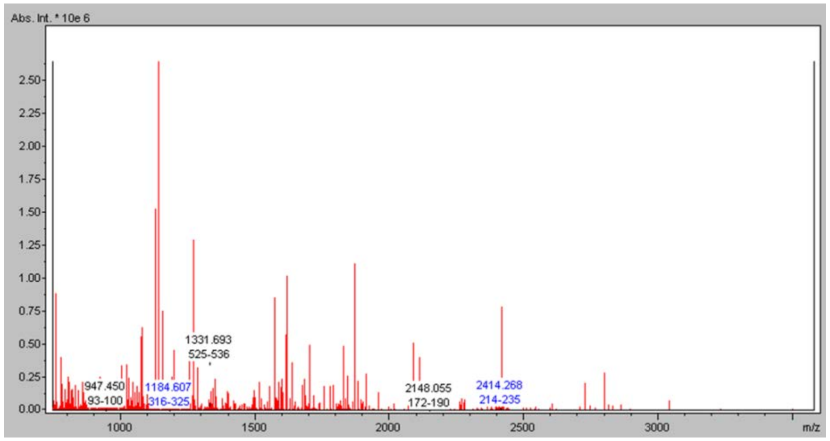


Tolerance (Da):0.500 Number of Peaks: 616

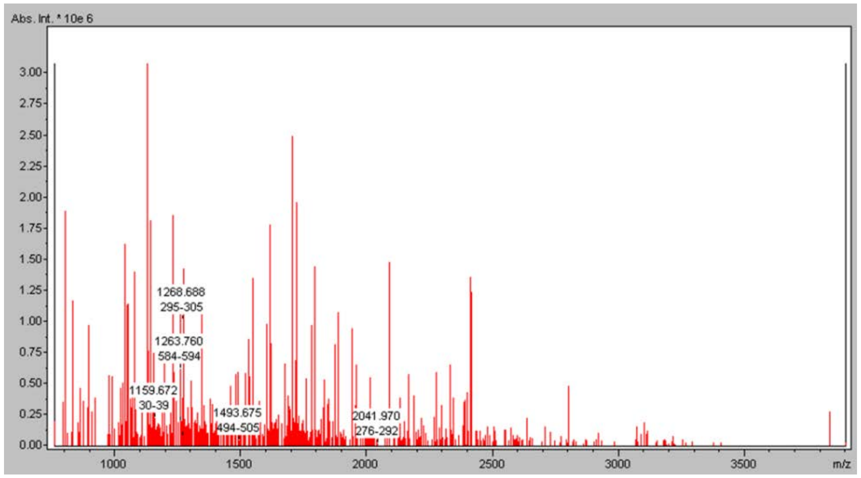


1 10 20 30 40 50 60 70
MAPIGLKAVV GEKIMHDVIK KVKKKGEWKV LVVDQLSMRM LSSCCKMTDI MTEGITIVED INKRREPLPS
80 90 100 110 120 130 140
LEAVYLITPS EKSVHSLISD FKDPPTAKYR AAHVFFTDSC PDALFNELVK SRAAKVIKTL TEINIAFLPY
150 160 170 180 190 200 210
ESQVYSLDSA DSFQSFYSPH KAQMKNPILE RLAEQIATLC ATLKEYPAVR YRGEYKDNAL LAQLIQDKLD
220 230 240 250 260 270 280
AYKADDPTMG EGPDKARSQL LILDRGFDPS SPVLHELTFQ AMSYDLLPIE NDVYKYETSG IGEARVKEVL
290 300 310 320 330 340 350
LDEDDDLWIA LRHKHIAEVS QEVTRSLKDF SSSKRMNTGE KTTMRDLSQM LKKMPQYQKE LSKYSTHLHL
360 370 380 390 400 410 420
AEDCMKHYQG TVDKLCRVEQ DLAMGTDAEG EKIKDPMRAI VPILLDANVS TYDKIRIILL YIFLKNGITE
430 440 450 460 470 480 490
ENLNKLIQHA QIPPEDSEII TNMAHLGVPI VTDSTLRRRS KPERKERISE QTYQLSRWTP IIKDIMEDTI
500 510 520 530 540 550 560
EDKLDTKHYP YISTRSSASF STTAVSARYG HWHKNKAPGE YRSGPRLIIF ILGGVSLNEM RCAYEVTQAN
570 580 590
GKWEVLIGST HILTPQKLLD TLKKLNKTDE EISS

**Fig. S4. Mass spectrometry identification data of the STXBP1 protein (see also Table S1) in three brain samples of six-month-old male rats.** The amino acid sequence of the STXBP1 protein is shown. The peptides identified by mass spectrometry are indicated in yellow.

FXR1 Fragile X mental retardation syndrome-related protein 1 OS=*Rattus norvegicus* GN=Fxr1 PE=1 SV=1 FXR1_RAT
MH+ (mono):1.008    MH+ (avg): 1.008

Tolerance (Da):0.500 Number of Peaks:775

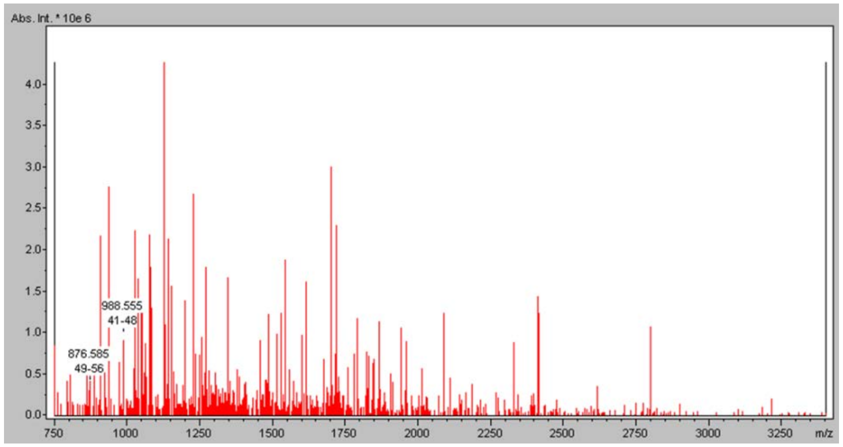


Tolerance (Da)  0.900   Number of Peaks: 1726

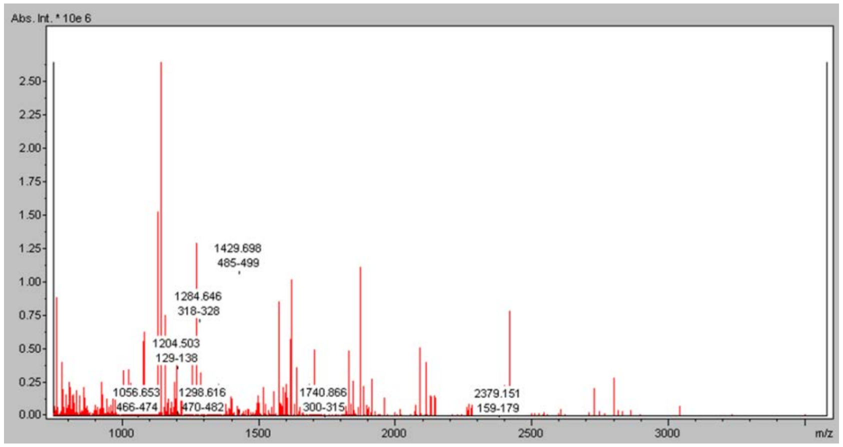


Tolerance (Da):0.500 Number of Peaks: 616

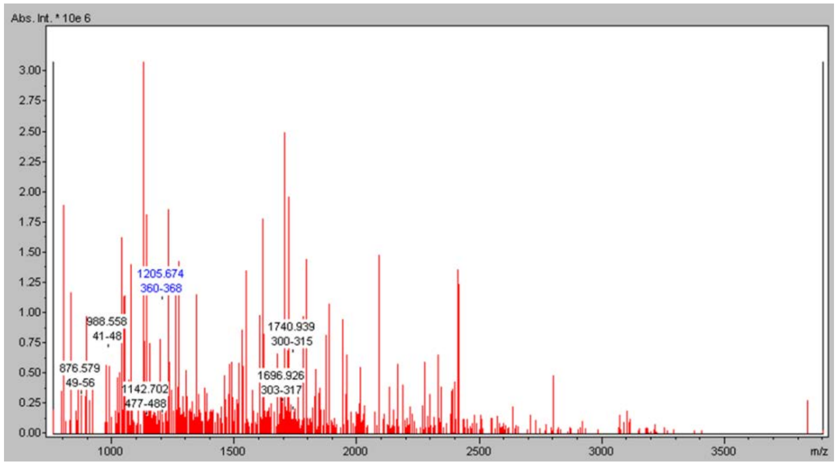


10 20 30 40 50 60 70
MAELTVEVRG SNGAFYKVFI KDVHEDSLTV VFENNWQPER QVPFNEVRLP PPPDIKKEIS EGDEVEVYSR
80 90 100 110 120 130 140
ANDQEPCGWW LAKVRMMKGE FYVIEYAACD ATYNEIVTFE RLRPVNQNKT VKKNTFFKCT VDVPEDLREA
150 160 170 180 190 200 210
CANENAHKDF KKAVGACRIF YHPETTQLMI LSASEATVKR VNILSDMHLR SIRTKLMLMS RNEEATKHLE
220 230 240 250 260 270 280
CTKQLAAAFH EEFVVREDLM GLAIGTHGSN IQQARKVPGV TAIELDEDTG TFRIYGESAE AVKKARGFLE
290 300 310 320 330 340 350
FVEDFIQVPR NLVGKVIGKN GKVIQEIVDK SGVVRVRIEG DNENKLPRED GMVPFVFVGT KESIGNVQVL
360 370 380 390 400 410 420
LEYHIAYLKE VEQLRMERLQ IDEQLRQIGM GFRPSSTRGP EKEKGYATDE STVSSVQGSR SYSGRGRGRR
430 440 450 460 470 480 490
GPNYTSGYGT NSELSNPSET ESERKDELSD WSLAGEDDRE TRHQRDSRRR PGGRGRSVSG GRGRGGPRGG
500 510 520 530 540 550 560
KSSISSVLKD PDSNPYSLLD NTESDQTADT DASESHHSTN RRRRSRRRRT DEDAVLMDGM TESDTASVNE
568
NGLGKRCD

**Fig. S5. Mass spectrometry identification data of the FXR1 protein (see also Table S1, Figure 1) in three brain samples of six-month-old male rats.** The amino acid sequence of the FXR1 protein is shown. The peptides identified by mass spectrometry are indicated in yellow.

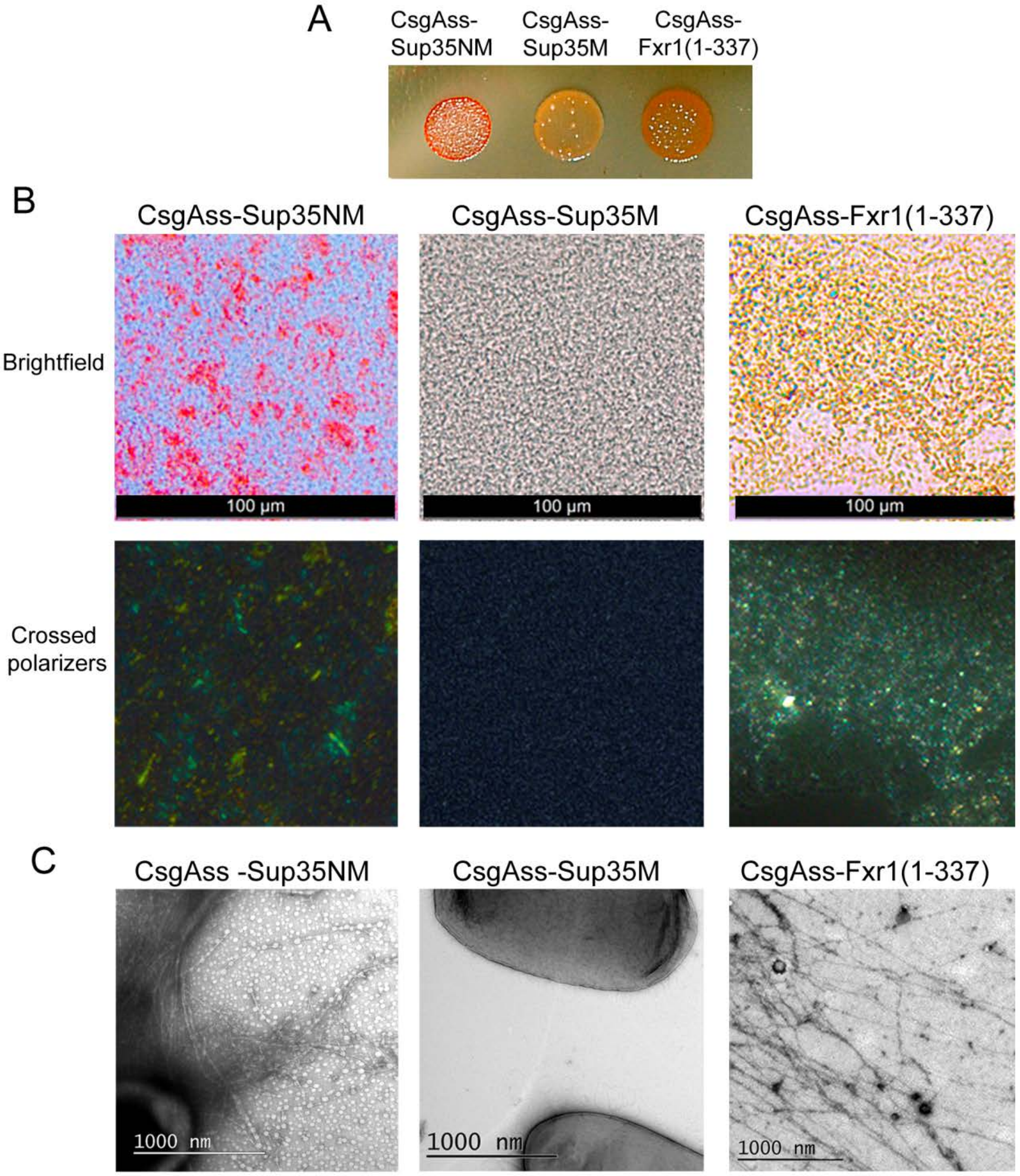


**Fig. S6.** FXR1 (1-337) fragment demonstrates amyloid properties in the bacteria-based C-DAG system.
(A) E.coli cells producing CsgAss-Sup35NM and CsgAss-FXR1(1-337) proteins form red colonies whereas cells producing CsgAss-Sup35M form pale colonies on agar plates containing CR.
(B) Micrographs of CsgAss-Sup35NM, CsgAss-Sup35M and CsgAss-FXR1(1-337) scraped cell samples harvested from CR-containing agar. Extracellular material containing CsgAss-Sup35NM and CsgAss-FXR1(1-337) binds CR (upper panel) and displays apple-green birefringence when viewed between crossed polarizers (lower panel). Scale bar, 100 μM
(C) Secreted CsgAss-Sup35NM and CsgAss-FXR1(1-337) proteins form fibrils visualized by transmission electron microscopy. Scale bar, 1000 nM.

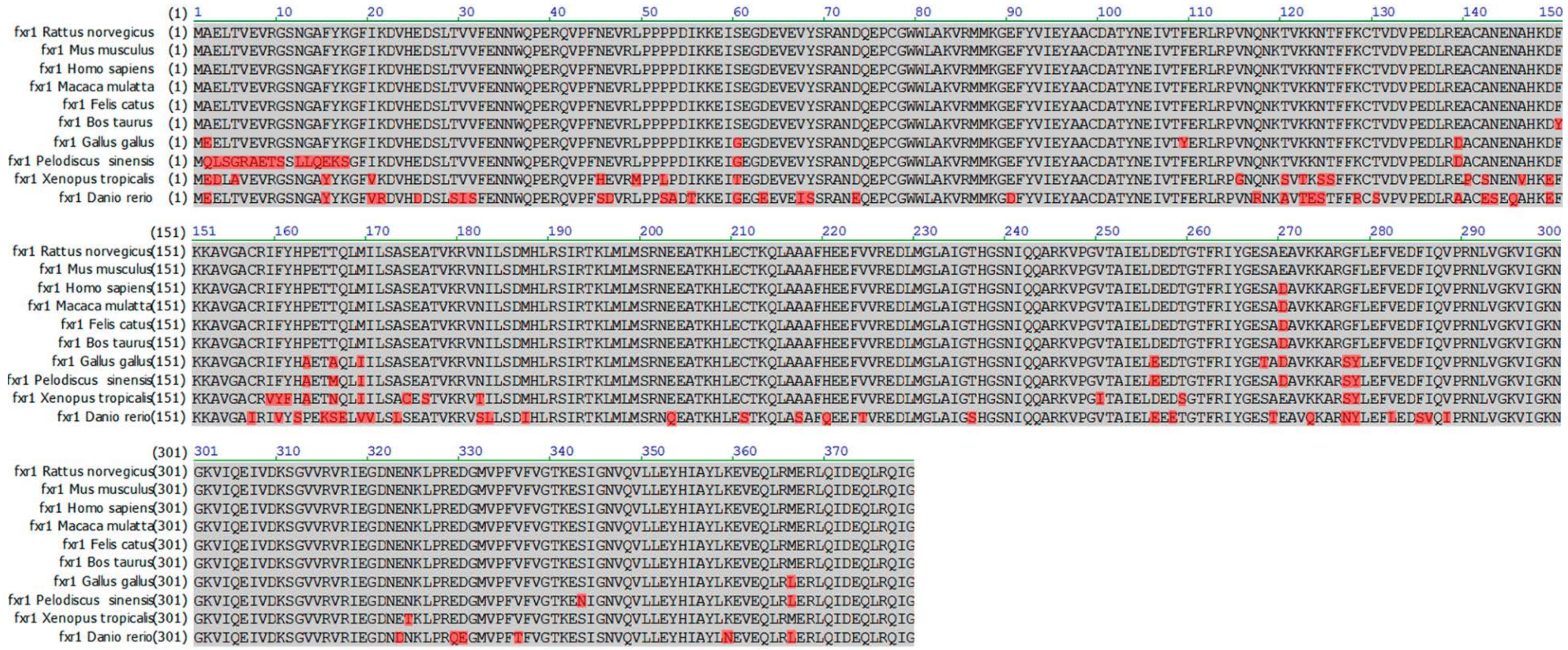

**Fig. S7. N-terminal amyloidogenic region of the FXR1 protein (1-379 aa) is highly conserved in mammals.** The amino acid sequence of the FXR1 protein of Wistar rats used in this work was identified and annotated in GenBank (accession number MG938503). This sequence was compared with corresponding canonical sequences of the FXR1 protein of different vertebrate species presented in UniProt database (http://www.uniprot.org). Sequence alignment for the FXR1 N-terminal domains of ten vertebrate species are presented. Residues matching those in rat sequence are shaded in grey, whereas amino acid substitutions are shaded in red.

**Supporting Table 1. The proteins forming SDS-resistant aggregates in rat brain**

| Protein | Function | Amyloidogenic regions of the protein as predicted by ArchCandy |
|---|---|---|
| NSF | Catalyzes the fusion of transport vesicles within the Golgi cisternae and is required for transport from the endoplasmic reticulum to the Golgi stack [4] | 118-145, 168-186, 350-374 |
| MBP | Has a role in the formation and stabilization of myelin layers [5] | - |
| RIMS1 | Involved in exocytosis, may act as scaffold protein that regulates neurotransmitter release at the active zone [6] | 758-780, 1538-1569 |
| STXB1 | Participates in the regulation of synaptic vesicle docking and fusion and is essential for neurotransmission [7] | 264-276, 405-430, 537-573 |
| FXR1 | RNA-binding protein [8], involved in regulation of long-term memory, mood and emotion [9,10] | 2-20, 225-247 |

**Supporting Table 2. Primers used in this work**

| Primer | Sequence |
|---|---|
| fxr1 EcoRI forward | atagaattcggcggagctgacggtgga |
| fxr1(379) BamHI reverse | ataggatccttacataccaatctgtcgc |
| fxr1 HindIII forward | tacaagcttatggcggagctgacggt |
| fxr1 (YFP)BamHI reverse | tatggatcccataccaatctgtcgcagctg |
| fxr1 (380) HindIII forward | attaagcttatgggtttcagaccttct |
| fxr1 (568) BamHI reverse | tacggatccatcacatcttttgcctagc |
| fxr1 NotI forward | atagcggccgcagagctgacggtggaggt |
| fxr1 XbaI reverse | gtattctagattaaccaatctgtcgcagct |
